# *In silico* identification of putative *Trypanosoma cruzi* enolase inhibitors

**DOI:** 10.1101/685206

**Authors:** Edward A. Valera-Vera, Melisa Sayé, Chantal Reigada, Mariana R. Miranda, Claudio A. Pereira

## Abstract

Enolase is a glycolytic enzyme that catalyzes the interconversion between 2-phosphoglycerate and phosphoenolpyruvate. In trypanosomatids enolase was proposed as a key enzyme after *in silico* and *in vivo* analysis and it was validated as a protein essential for the survival of the parasite. Therefore, enolase constitutes an interesting enzyme target for the identification of drugs against Chagas disease. In this work, a combined virtual screening strategy was implemented, employing similarity virtual screening, molecular docking and molecular dynamics. First, two known enolase inhibitors and the enzyme substrates were used as queries for the similarity screening on the Sweetlead database using five different algorithms. Compounds retrieved in the top 10 of at least three search algorithms were selected for further analysis, resulting in six compounds of medical use (etidronate, pamidronate, fosfomycin, acetohydroximate, triclofos, and aminohydroxybutyrate). Molecular docking simulations predicted acetohydroxamate and triclofos would not bind to the active site of the enzyme, and a re-scoring of the obtained poses signaled fosfomycin and aminohydroxybutyrate as bad enzyme binders. Docking poses obtained for etidronate, pamidronate, and PEP, were used for molecular dynamics calculations to describe their mode of binding. From the obtained results, we propose etidronate as a possible *Tc*ENO inhibitor, and describe desirable and undesirable molecular motifs to be taken into account in the repurposing or design of drugs aiming this enzyme active site.

## 1. Introduction

Chagas disease, also known as American trypanosomiasis, is a neglected tropical infection caused by the hemoflagellate protozoan parasite *Trypanosoma cruzi* [1]. According to the World Health Organization, around 6-7 million people worldwide are estimated to be infected with this parasite, mostly in 21 Latin American countries [2]. Moreover, Chagas disease is a global health problem mainly because of migration of people from endemic countries to the rest of the world [3]. Benznidazole and nifurtimox are the only drugs approved for the treatment of Chagas disease, which induce serious side-effects and have very limited efficacy, especially in the chronic phase of the disease [4]. Thus, studies focusing on the development of new therapeutic alternatives and on the identification of novel drug targets are still necessary.

Enolase (2-phospho-D-glycerate hydrolase, EC 4.2.1.11) is a key enzyme of the processes of glycolysis and gluconeogenesis and it catalyzes the reversible dehydration of 2-phosphoglycerate (2-PG) to phosphoenolpyruvate (PEP) in the presence of the metal ion magnesium (Mg^2+^). Enolase is found in a wide variety of organisms ranging from bacteria to mammals including trypanosomatids [5]. Peptide mass fingerprinting allowed the detection of this enzyme in the different stages of *T. cruzi* life cycle, with a higher expression in trypomastigotes and amastigotes (forms present in the mammalian host) compared to epimastigotes (form present in the insect vector) [6]. Glycolysis and gluconeogenesis play crucial roles in the ATP supply and synthesis of glycoconjugates which are important for the viability and virulence of the human-pathogenic stages of *T. cruzi, T. brucei* and *Leishmania* spp. [7], which are the etiological agents of the most important neglected diseases around the world. In addition to its metabolic importance, enolase has been proposed as a virulence factor in *Leishmania* spp. because of the enzyme ability to bind plasminogen [8], Therefore, enolase constitutes an interesting enzyme target for the identification of drugs for Chagas’ disease and also to other trypanosomatid-caused diseases.

Phosphonoacetohydroxamate (PAH) is a potent enolase inhibitor which has nanomolar values of IC_50_ inhibitory activity against this enzyme from different species [9, 10]. The crystal structure of the *T. brucei* enolase (*Tb*ENO) with this inhibitor has already been determined, showing that PAH and the natural substrate PEP bind to the same residues in the structure [11, 12]. Recently, another highly potent enolase inhibitor, called SF2312 has been reported [13]. SF2312 is a phosphonate antibiotic of unknown mode of action produced by the actinomycete *Micromonospora*. Moreover, *Tb*ENO has been validated as a drug target by RNA interference, which led to a negative effect on bloodstream trypomastigotes growth within 24 h and produced the death at 48 h. The same study also demonstrated that 80-85% reduction on enolase activity was enough for cell death to occur [14]. This evidence suggests that incomplete inhibition of this enzyme *in vivo* might prove sufficient for effective treatment.

Considering the relevance of enolase in trypanosomatids, in this work, we applied a combined virtual screening strategy, based on known ligands (2-PG, PEP, PAH and SF2312) and the protein target enolase, in order to find novel *Trypanosoma cruzi* enolase (*Tc*ENO) inhibitors from human-approved drug databases.

## 2. Methodology

### 2.1. Identification of the target protein

A screening in the TDR “Targets Database”(v5) (http://tdrtargets.org/) was performed to obtain information about a potential drug target for Chagas disease [15]. The screening was restricted to *T. cruzi* genes coding for enzymes that produce a significant loss of fitness when absent or inhibited in the parasite or in related organisms. Resulting targets were then evaluated by bibliographical searching, and the *Tc*ENO was selected for identification of possible binders to the active site, as well as sequence and structural comparison with its homologues in humans, *Trypanosoma brucei*, and *Leishmania Mexicana.* Amino acid sequence analysis were performed using the T-Coffee multiple sequence alignment package [16] and structural alignment on PyMOL [17].

### 2.2. Ligand-based virtual screening

Computational approaches for the identification of likely *Tc*ENO inhibitors started with a similarity-based search with molecules that experimentally bind to the active site of the protein, the enzyme substrates (PEP and 2-PG) and two previously characterized inhibitors (PAH and SF2312). The screening was performed on the Sweetlead database, that comprises about 10,000 drugs and isolates from traditional medicinal herbs [18], using the software LiSiCA v1.0 (Ligand Similarity using Clique Algorithm) [19], KCOMBU (K(ch)emical structure COMparison using the Build-up algorithm) [20], and ShaEP (Molecular Overlay Based on Shape and Electrostatic 5Potential) v 1.1.2 [21]. The structures of the database were ordered with five different criteria: according to (I) the 2D and (II) 3D similarity using the LiSiCA software based on the Tanimoto similarity index [22], (III) using the Ikcombu algorithm provided by the KCOMBU software, (IV) by shape and (V) electrostatic potential employing the ShaEP software. These algorithms were selected on the basis of the results obtained after doing the search with some known molecules, after discarding other algorithm that did not show the query molecule in the first position of the obtained lists. Following, for each of the characterized ligands, compounds that appeared between the first ten in at least three of the five ranking criteria were selected for further analysis.

### 2.3. Target-based virtual screening

The structural files of the molecules selected by similarity screening were then obtained from the ZINC database [23]. Preparation of the PDBQT files (Protein Data Bank, partial charge (Q), and atom type (T)) was performed using AutoDock Tools v1.5.6 [24].

A search in the Protein Data Bank (PDB) [25] was carried to find the crystal structure of *Tc*ENO (PDB: 4G7F); but for being in apo conformation and lacking the residues ASTG(39-42), which are involved in the interaction with the phosphate groups of the ligands PEP and PAH, a structural model of the closed conformation was predicted by fragment assembly simulation using the server I-TASSER [26]. The model was validated by calculating its DOPE and GA414 scores with Modeller [27], generating a Ramachandran plot in the online server Rampage [28], and by redocking of PEP and PAH to see if the docking software was able to reproduce the poses of the ligands co-crystalized with the *Tb*ENO.

For molecular docking assays we used AutoDock 4.5 [29], with a grid map of 50 x 58 x 44 points, spacing of 0.0375 nm, and centered on the position occupied by PEP in the *Tb*ENO crystal structure, surrounding the residues conforming the enzyme active site (Ala39, Ser40, Lys155, His156, Glu165, Asp243, Glu291, Asp318, Lys343, His371, Arg372, Ser373, and Lys394).

As enolase needs two dicationic ions in the active site to bind its substrates, magnesium ions were added to the PDBQT file of the receptor, according the coordinates of the magnesium ions found in the 2PTY pdb file. For each ligand we did 100 docking runs with the Lamarckian Genetic Algorithm, with a population size of 300, and 2.7х10^4^ as the maximum number of generations, followed by conformational clustering of the obtained poses into groups with RMSD <0.2 nm. All the obtained docking poses were visually analyzed using PyMOL [17], and rescored using the neural-network based scoring software NNScore v1 [30].

### 2.4. Pharmacophore modeling

Structural alignments and pharmacophore modeling were performed using LigandScout 4.1.10 (Inte:Ligand, Software-Entwicklungs und Consulting GmbH, Austria) [31], which license was kindly granted to our laboratory by the developer company. Modeling was performed using the LigandScout default parameters, with the structures of the four reported ligands (PEP, 2-PG, PAH, and SF2312) and the molecules predicted to interact with the binding site in the docking assays (etidronate and pamidronate). The model obtained shows H-bonds acceptor/donors and ionizable atoms expected for a molecule that is able to bind the active site of the enzyme. The complete pipeline of the screening strategy is summarized in Fig.1.

**Fig. 1.**
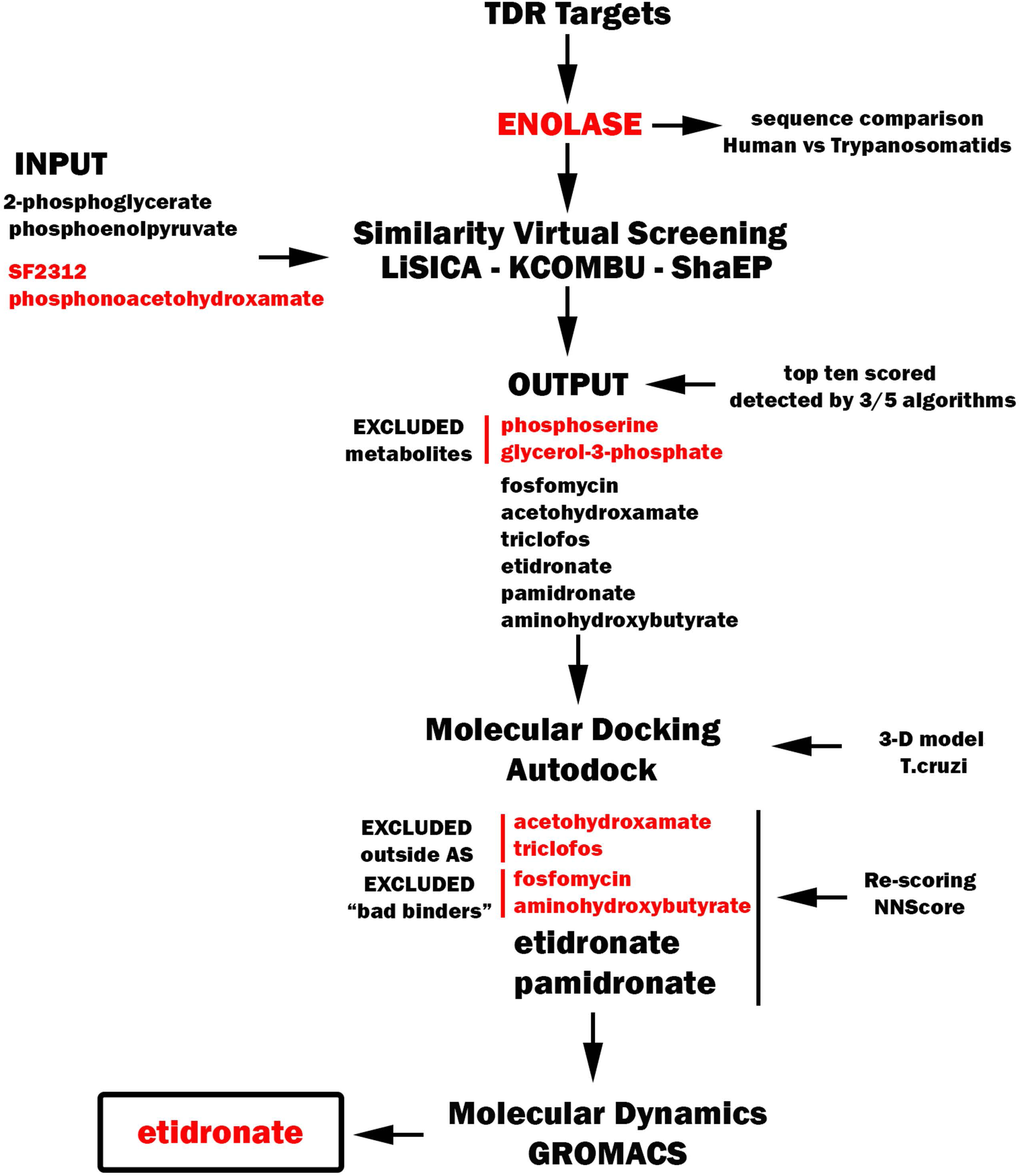
Pipeline of the screening strategy. A two-steps methodology was applied for identification of structure candidates to bind the *Tc*ENO active site. The ligand-based virtual screening was performed using natural substrates (2-PG and PEP) and known inhibitors (PAH and SF2312) as reference molecules, followed by a receptor-based strategy using the modeled *Tc*ENO as the receptor and the compounds obtained from the similarity screening as ligands. Docking poses were subsequently evaluated by re-scoring and molecular dynamics simulations

### 2.5. Molecular Dynamics

The poses with less free energy obtained from AutoDock 4.5 for PEP, etidronate and pamidronate were assessed by molecular dynamic (MD) simulations using GROMACS 5.0.5 [32]. For this, the topology of the *Tc*ENO (including the magnesium ions) was calculated using the GROMOS96 43a1 force field. The topology of the *Tc*ENO in complex with the ligands was obtained by merging the topology obtained for each ligand from the PRODRG server [33], and embedded in the *Tc*ENO topology file. Water molecules and ions were added to each *Tc*ENO-ligand system in a dodecahedral solvation box, using the SCP water model. After energy minimization of the system, restraints were applied to the position of the protein and the ligand. The system was then heated to 300 K at constant volume (NVT) during 50 ps, using a modified Berendsen thermostat with a temperature coupling time constant (Tt) of 0.1 ps; followed by 50 ps of pressure equilibration (NPT) with isotropic Berendsen barostat at 1.0 bar of pressure at a time constant (Tp) of 2.0 ps. Restraints were removed and the MD production run was extended for 50 ns at a pace of 2 fs, recorded every 10 ps for further analysis.

Periodic boundaries conditions were removed from the resulting trajectories to further align them using the RMSD visualizing tool on VMD [34], fitting the trajectories to the backbone of the protein. The snapshots were then visually analyzed and RMSD was calculated for the protein and the ligands. On VMD, the H-bonds formed across the trajectories between the ligands and the protein were obtained for all the trajectories, setting a 0.35 nm bond distance and angle of 30°. The residues with a distance to the ligand <0.5 nm were considered for contact counting with ‘gmx mindist’, and interaction energy calculations with ‘gmx energy’. The binding free energies of the simulated complexes were calculated with the tool GMXPBSA 2.1 [35] on 100 frames extracted from the last 30 ns of simulation. Principal component analysis (PCA) was performed on the same time lapse of the trajectories, using the gromacs tools ‘gmx covar’ and ‘gmx anaeig’, followed by a free energy landscape (FEL) generated for the projection of the first two principal components (PC1 and PC2) using ‘gmx sham’ and ‘xpm2ps’.

## 3. Results and Discussion

### 3.1. Target analysis

A screening restricted to *T. cruzi* genes coding for enzymes which inhibition or deletion also produce a significant loss of fitness in bloodstream forms of *T. brucei* (no information is available for *T. cruzi*) using the TDR “Targets Database” (v5) to retrieve potential drug targets was performed. The enzyme *Tc*ENO was present among the group of *T. cruzi* genes resulting from the screening. In *T. brucei,* the knock-out of this this enzyme also produced a validated loss of fitness in the bloodstream and procyclic forms of the parasite, in addition to affect the differentiation between the procyclic to bloodstream forms. The enolase is also indicated as a known druggable target in humans in its three isoforms. A multiple alignment between the *Tc*ENO, *Tb*ENO, *Lm*ENO, and the three human enolases (*Hs*ENO1, 2, and 3) shows that residue Lys155, conserved in the enzyme sequence of all three trypanosomatids evaluated, is replaced by a serine in the human homologues (Online Resource 1). This residue belongs to a very flexible loop on the active site, being able to move towards the cavity or accommodating away from it [36], altogether pointing the exploitability of this residue in the design of drugs specific to trypanosomatid enolases. These results indicate that *Tc*ENO meets the requirements to be an excellent drug target: it is essential in all stages of the parasite, including the processes of differentiation between them; it is druggable, as other enolases have known inhibitors; it has differences with the human homologue; and its enzymatic activity is easy to measure.

### 3.2. Similarity screening

The first step of the screening for putative *Tc*ENO inhibitors was a ligand-based strategy using the enzyme substrates (2-PG and PEP, Fig.2a) and previously characterized inhibitors (PAH and SF2312, Fig.2b) as query structures. The Sweetlead database was screened using three different similarity search software with a total of five different algorithms, as detailed under “Methodology”. For each query molecule, the top ten scored compounds that were detected at least by three algorithms were chosen for further analysis. Using these criteria eight possible inhibitors of the *Tc*ENO were obtained; six of them of medical use and two human metabolites. These compounds were: I) etidronate and II) pamidronate, two bisphosphonates used to treat Paget’s disease of bone and to prevent and treat heterotopic ossification; III) fosfomycin, an antibiotic used to treat infections of the urinary tract; IV) acetohydroxamate, used to lower the level of ammonia in urine in some types of urinary infections; V) triclofos, a sedative drug used for treating insomnia, VI) aminohydroxybutyrate, an anticonvulsant which is used for the treatment of epilepsy, and the human metabolites VII) phosphoserine and VIII) glycerol-3-phosphate (Fig.2c). The two metabolites were excluded from further evaluation.

**Fig. 2.**
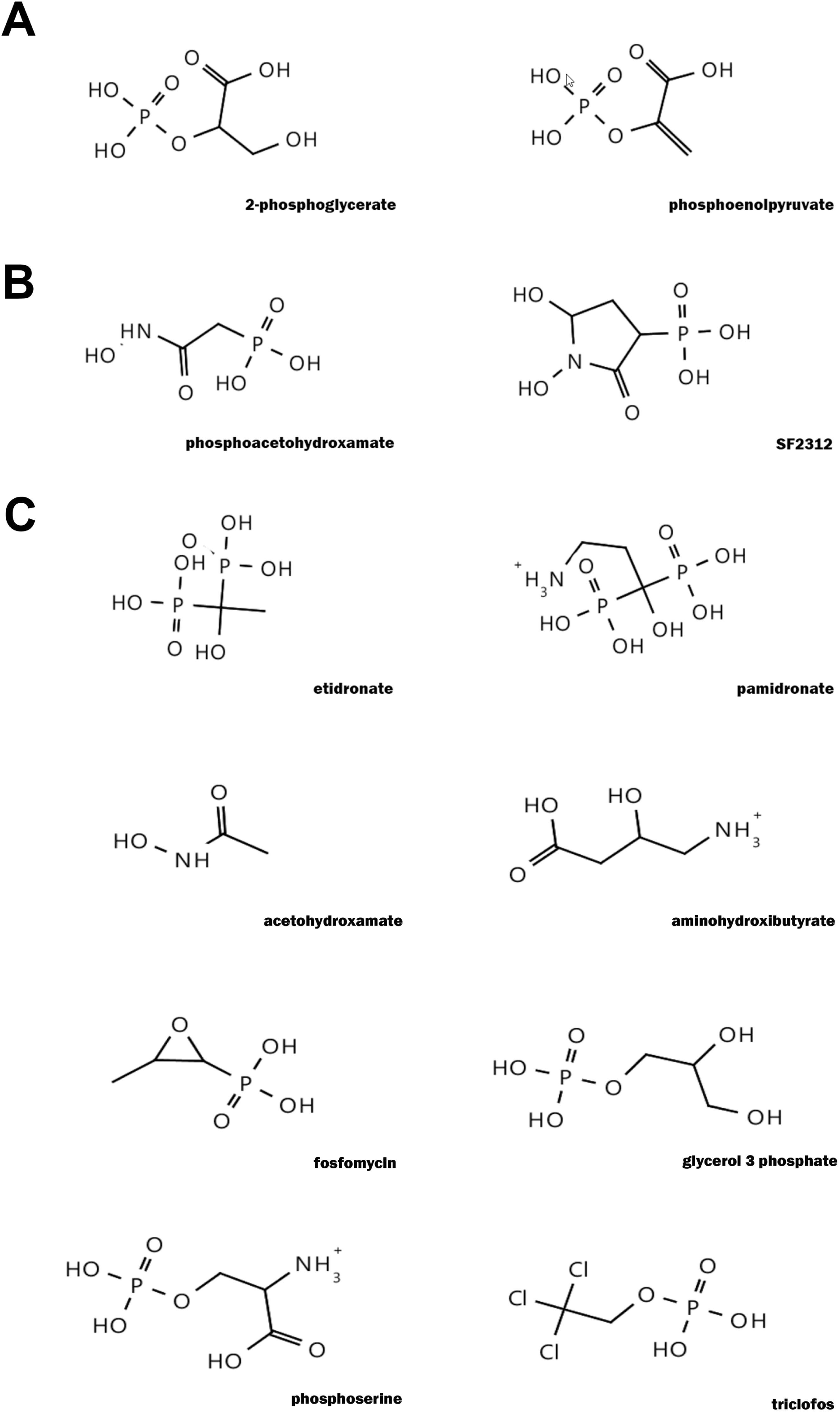
Chemical structures of a) natural substrates, b) known inhibitors and c) putative binders of the *Tc*ENO obtained through similarity screening as described under “Materials and Methods”

### 3.3. *Tc*ENO structure modeling

In the production of the 3D model for the *Tc*ENO, the I-TASSER simulations converged in a single structure, which signals a good quality of the model. To further assessing the quality of the model we used various approaches. The quality score that I-TASSER gives for the models it produces, the C-score, was 1.77 for the *Tc*ENO; this score ranges from −5 to 2, where the best score being 2. The Ramachandran plot shows only a residue (Arg400) falling in the outlier region, with every other residue inside the favored or allowed regions (Online Resource 2). The GA341 and the DOPE scores calculated with Modeller were respectively 1.0 and - 15.55; the later ranges from 0.0 to 1.0, where values nearest or equal to 1.0 signal a good quality, while the former represents the energy of the protein, where lower values are assigned to better quality models. Altogether, these metrics show that the predicted structure has an overall good quality.

### 3.4. Molecular docking

First, we assessed the docking model by ‘redocking’ the ligands PEP and PAH with the *Tc*ENO model and comparing the docked poses with that of the crystalized ligands in the homologous *Tb*ENO (PDB: 2PTY and 2PTZ) [11] by calculating their RMSD. Initially we were not able to reproduce the crystalized poses, as the enolase requires two dicationic ions in the active site; so, we decided to include two magnesium ions in the same coordinates as in the *Tb*ENO crystals. Because ions interacting with a protein will have a charge different than the ion in solution, and improper manage of such charges would result in incorrect docking poses [37], we tested docking with different charges of the ions in the range from +2 to +0.5, in order to find the charge values which produces the poses closest to the crystalized ligands. After these changes, AutoDock 4.5 achieved to reproduce the crystalized poses in a single cluster with a RMSD of 0.17 nm compared to the crystals when both magnesium had a charge of +0.8, which was maintained for all the ligand docking calculations.

From the six probed ligands, acetohydroxamate and triclofos did not docked in the active site of the enzyme in any of the generated poses. A re-scoring in NNScore of the poses for the four remaining compounds labeled fosfomycin and aminohydroxybutirate as “bad binders”, being left etidronate and pamidronate as promising candidates for binding in the active site of *Tc*ENO. Fig.3 shows the generated poses for etidronate, pamidronate, PAH, and PEP, with the *Tc*ENO residues predicted to form h-bonds with each ligand. For PEP and PAH all the predicted h-bonds are listed in the PDB entries for their complexes with *Tb*ENO. All four compounds share a phosphate motif, which appears interacting in the same manner with the residues Ala39, Ser40, Arg372, Ser373, and one of the magnesium ions, pointing the importance of these interactions for the binding mode, suggesting any future drug designed to inhibit *Tc*ENO should retain highly electronegative atoms in this position. The remaining phosphate in etidronate and pamidronate is shown interacting with Arg372, Lys3443, Lys394, and the two magnesium ions, in a similar fashion as the carboxyl group in PEP, and in PAH the hydroxyl and the carbonyl groups, implying the necessity of electronegative groups in that space for the ligands to bind. The hydroxyl group in etidronate in was located towards the magnesium ions and residue Gln164, Glu165, and Glu208; while the methyl group was pointing towards Lys343 and Arg372. For pamidronate, the pose was similar to that of etidronate, with the distinct amine in the direction of Lys343, Gln165, and Glu208; extending towards His371.

**Fig. 3.**
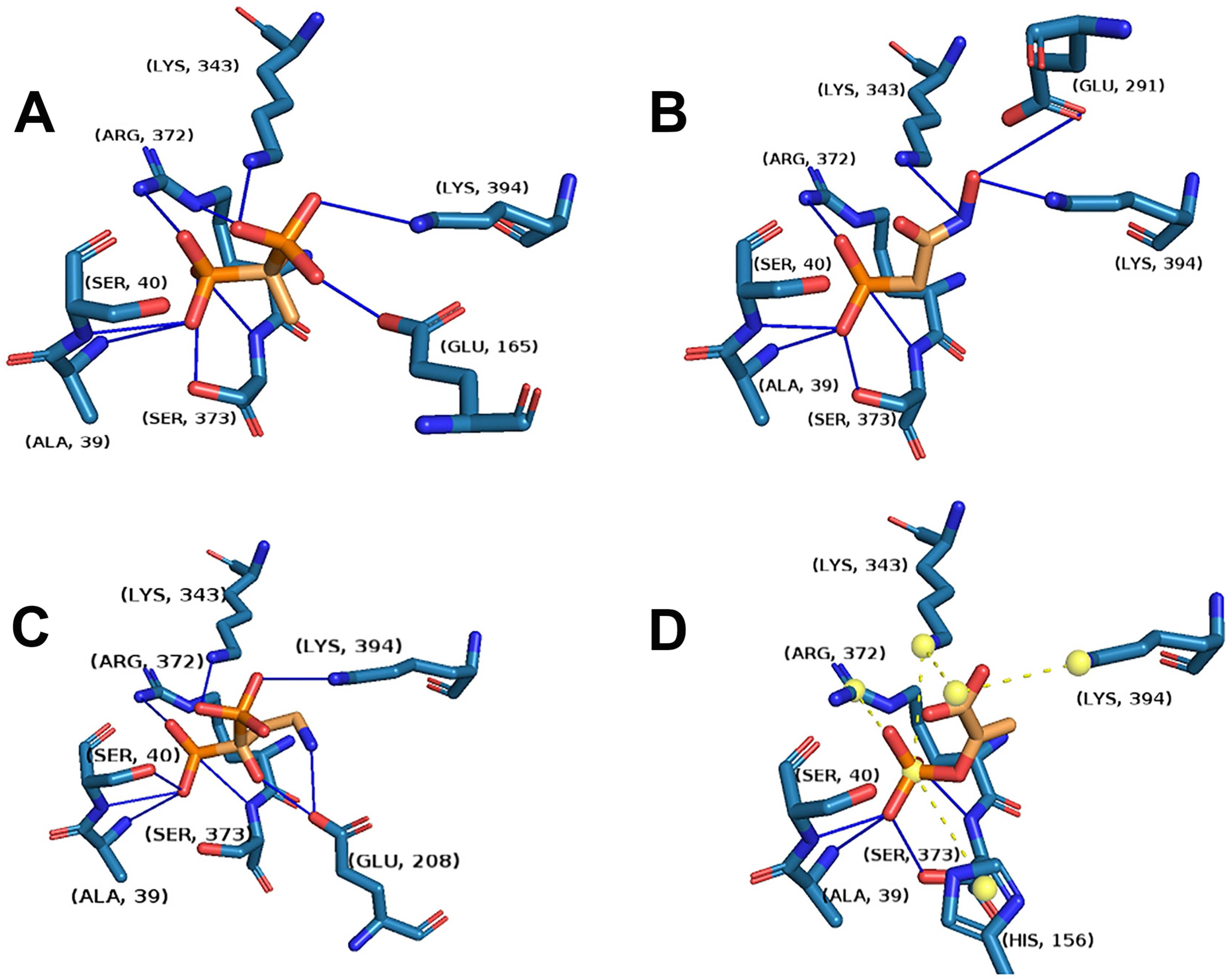
Molecular docking poses of a) etidronate, b) PAH, c) pamidronate, d) PEP, showing the residues that bind to the compounds (orange). Each pose corresponds to the lowest score obtained with each docked ligand. Hydrogen bonds are indicated as blue lines, salt bridges as yellow dotted lines, and charges centers as yellow spheres

### 3.5. Pharmacophore model

A pharmacophore was constructed using the structures that bind into the active site of the enolase experimentally (PEP, 2-PG, PAH, SF2312), and those predicted to be good binders (etidronate and pamidronate) (Online Resource 3a). Both enzyme natural substrates present a negative ionizable phosphate group capable of accepting hydrogen bonds, and also other hydroxyl or carboxyl residues that are capable of H-bonding (Online Resource 3c-d). The known inhibitors, PAH and SF2312, also share these critical chemical structural features, containing a negative ionizable and H-bond acceptor hydroxamate functional group in addition to phosphonate groups that can form H-bond as well (Online Resource 3c-d). Accordingly, the expected molecules containing the pharmacophore might have H-bond acceptor groups such as phosphonates, ether and/or hydroxyl groups. The phosphonate groups could also fulfill the pharmacophoric requirements of negative ionizable groups located at the indicated positions (Online Resource 3b). As can be seen in Online Resource 3c, all the structures share the five H-bond acceptors and the negative ionizable group except aminohydroxybutyrate and fosfomycin. All mentioned compounds with the H-bond acceptor and non-ionizable groups marked are shown in Online Resource 3d.

### 3.6. Evaluation of docking poses

In order to describe in more detail the possible mode of interaction of etidronate and pamidronate with the protein, as well as to address the flexibility of the receptor in the interaction, we performed MD simulations on the apo form of the protein, and in complex with both ligands.

#### 3.6.1. Root Mean Squared Deviation

Simulations of 100 ns were run with Gromacs for *Tc*ENO in apo form. The backbone RMSD for was lower than 0.3 nm, achieving stability after 10 ns, so for the complexes with ligands we did the simulations for 50 ns.

MD for the protein complex with PEP, etidronate, and pamidronate resulting from the molecular docking also had backbone RMSD values (Fig.4a) lower than 0.3 nm. For PEP the mean value between 20 and 50 ns was lower than the apo form, as expected for a protein stabilized by interactions with its ligand. For etidronate the value was even lower, which could mean the interactions with this ligand allows for less conformational changes of the protein during the simulation. In the case of the pamidronate complex the backbone RMSD is similar to that of PEP up to 25 ns, when it increases reaching values similar to that of the apo form, suggesting an increase in the protein flexibility after this point in the simulation.

**Fig. 4.**
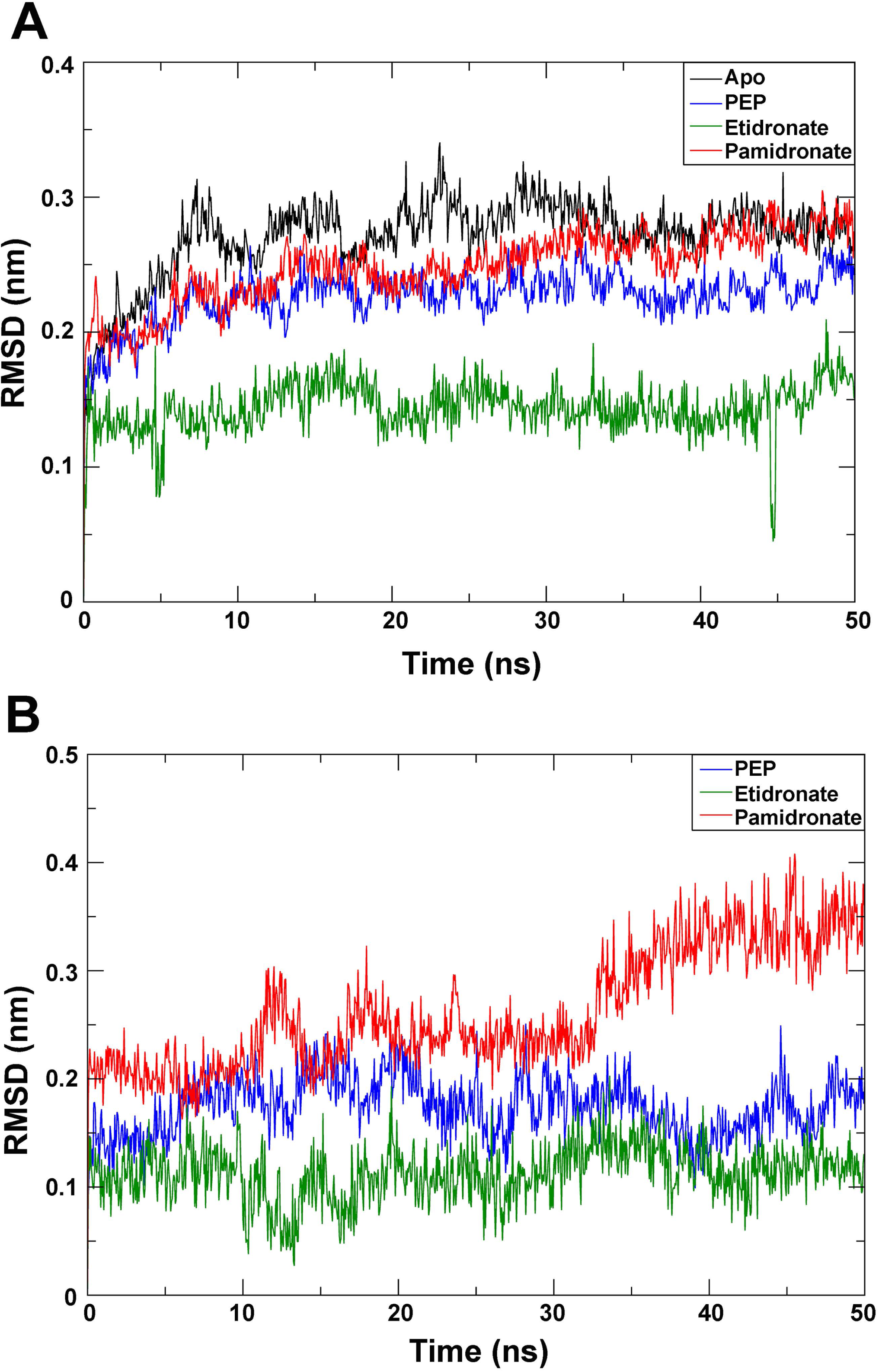
RMSD plot of *Tc*ENO backbone (a) and the probed ligands (b) during the MD simulation

RMSD of the ligands (Fig.4b) are lower than 0.2 nm for PEP and etidronate, and show less variation for etidronate than PEP, suggesting again the interactions between the former and *Tc*ENO allow for less flexibility than for the former. For pamidronate the ligand RMSD was higher than for the other two, and it shows an increase after 30 ns to values higher than 0.3 nm, indicating big conformational changes that in addition to the increase in flexibility of the protein might be the product of the loss of the interactions predicted by the molecular docking. This was confirmed upon visual examination of the frames, where pamidronate starts with both phosphate groups interacting in the same pocket as etidronate and the amine wobbling, followed by a motion of the whole molecule towards a more distal position in the pocket.

#### 3.6.2. H-bonds

Fig.5 shows the h-bonds that appeared during the simulation in at least 10% of the analyzed frames. Between PEP and *Tc*ENO the residue Ser40 formed the most stable H-bonds with the phosphate group of PEP; one of them lasting for 93% of the simulation involving the OH group from the side chain, and the other with the NH from the backbone in 85% of the trajectory. The third most stable H-bond was between the carboxyl of the backbone in Ala39 and the phosphate group, occurring in 45% of the analyzed snapshots. Two more H-bonds took place during 30% of the simulation involving the carboxyl group in PEP, with the side chains of Gln164 and Lys343. All the other H-bonds found occurred in less than the 10% of the simulation; involving residues Arg15, His44, Lys155, His156, Ser370, Arg372, Ser373, and Lys394.

**Fig. 5.**
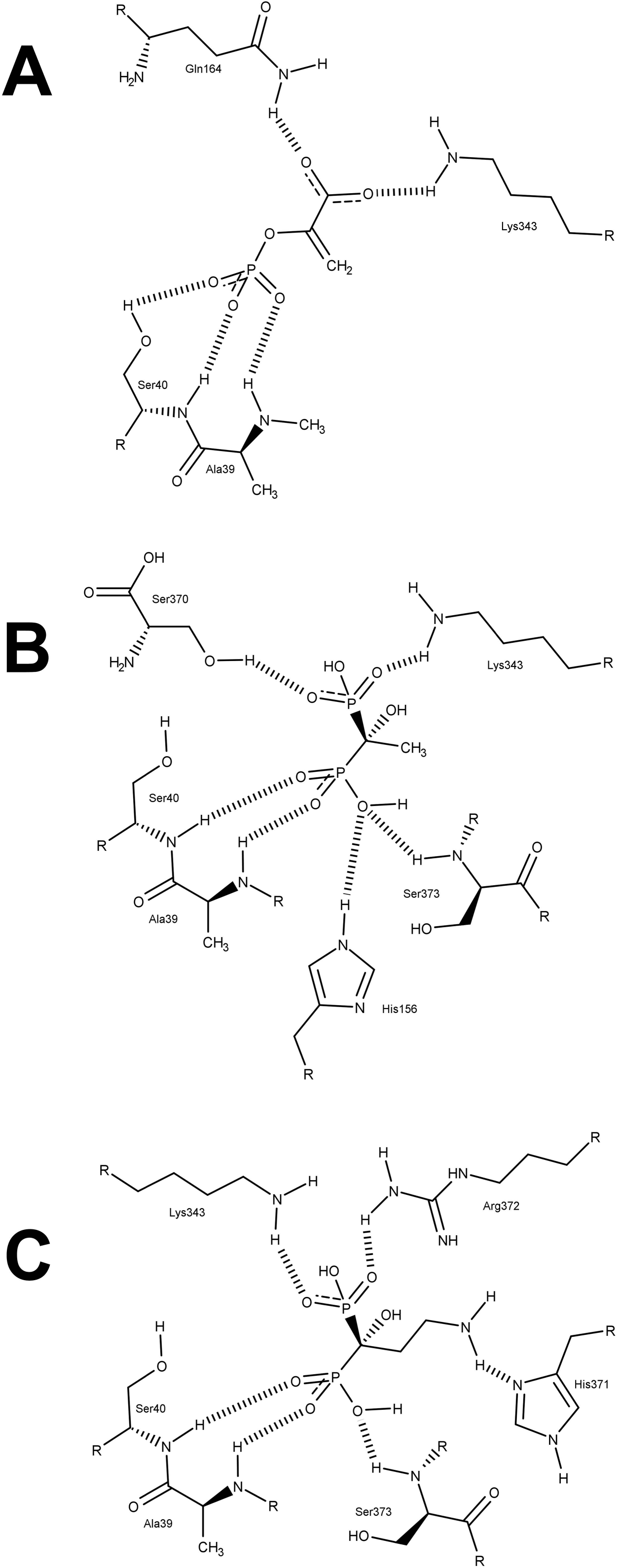
2D representation of the H-bonds formed in at least 10% of the analyzed MD frames between amino acid residues in *Tc*ENO and (a) PEP, (b) etidronate, and (c) pamidronate

For the complex *Tc*ENO-etidronate one of the phosphate groups in the ligand established H-bonds with the NH in the main chains of Ala39 and Ser40 during 81% and 73% of the frames respectively. The same phosphate received H-bonds from the NH in the backbone of Ser373 in 58% of the frames, and the side chain NH in His156 18% of the time. The most recurring H-bonds with the second phosphate of etidronate involved the side chains of Ser370 and Lys343, establishing such interactions during 32% and 30% of the simulation, respectively. Other H-bonds appearing in less than 10% of the snapshots involved Gln16, Ala39, Ser40, Arg372, Ser373, and Lys394.

In the case of pamidronate H-bonds were less frequent, with the backbone NH of residues Ala39, Ser40 and Ser373 establishing such interactions with the ligand in 38%, 14% and 12% of the frames, respectively. His371 and Arg372 side chains had H-bonds with pamidronate in 11% of the simulation, and many other residues establishing H-bonds in less than 10% of the run.

#### 3.6.3. Binding energy and residue interaction

Binding free energy of the complexes with the ligands was calculated with GMXPBSA 2.0 (Table 1). *Tc*ENO-PEP complex had the lowest binding ΔG, with a predominant electrostatic contribution. For *Tc*ENO-etidronate the binding energy was 2.5 times higher but still negative, while in *Tc*ENO-pamidronate the binding energy was around zero, as a result highly positive polar contribution, which can be attributed to the NH_2_ group for being the only polar substituent in pamidronate that is not present in etidronate. The low binding energy for the pamidronate is consistent with the increase in RMSD of the ligand and its translocation inside the active site, signaling the loss of the interactions predicted by the molecular docking.

**Table 1.** Binding free energies calculated with MMPBSA (kcal/mol) for PEP, etidronate, and pamidronate interacting with the active site of *Tc*ENO

| Ligand | PEP | Etidronate | Pamidronate |
| --- | --- | --- | --- |
| Coulombic | -3593.70 ± 12.72 | -1305.35 ± 24.53 | -399.54 ± 4.01 |
| Lennard-Jones | 111.98 ± 16.77 | -111.13 ± 17.68 | -114.09 ± 12.21 |
| Polar solvation | 5.15 ± 1.08 | 10.67 ± 1.54 | 514.36 ± 3.30 |
| Non-polar solvation | -7.73 ± 0.03 | -8.20 ± 0.02 | -10.71 ± 0.02 |
| Polar contribution | -3588.5 | -1294.7 | 114.8 |
| Non-polar contribution | 104.2 | -119.3 | -124.8 |
| Binding free energy | -3484.30 ± 19.05 | -1414.01 ± 28.02 | -9.98 ± 12.63 |

To address the individual contribution of residues to the interaction we summed the Coulombic and Lennard-Jones energies that were in contact with the ligands in more than 50% of the analyzed frames. As listed in Table 2, the residues with the lowest interaction energy with PEP are all noted in the PDB entry of 2PTY as the involved on the binding, which validates the capability of the model to predict the experimental binding mode. Regarding the bisphosphonates, the interaction energies with pamidronate are lower compared to etidronate, and both of them lower than PEP.

**Table 2.** Percentage of frames with contacts and interaction energy (IE) between the ligands and *Tc*ENO amino acid residues with more than 50% of contacts

| Ligand | Residue | % contacts | Coulomb | LJ | IE |
| --- | --- | --- | --- | --- | --- |
| PEP | 38 | 97 | -3.7 | -2.4 | -6.2 |
|  | 39 | 100 | -28.5 | -5.3 | -33.8 |
|  | 40 | 100 | -100.7 | 7.8 | -92.9 |
|  | 44 | 70 | -13.9 | -6.0 | -19.9 |
|  | 155 | 68 | -26.8 | -2.2 | -28.9 |
|  | 156 | 76 | -1.6 | -0.8 | -2.4 |
|  | 164 | 100 | -27.5 | -6.7 | -34.2 |
|  | 245 | 100 | -0.3 | -3.4 | -3.8 |
|  | 341 | 100 | 0.0 | -3.7 | -3.6 |
|  | 343 | 100 | -55.7 | -2.4 | -58.1 |
|  | 370 | 97 | -10.2 | -0.8 | -11.0 |
|  | 372 | 100 | -15.2 | -3.2 | -18.4 |
|  | 394 | 100 | -41.2 | 0.2 | -41.0 |
| Etidronate | 38 | 100 | -7.1 | -5.4 | -12.5 |
|  | 39 | 100 | -40.3 | -5.8 | -46.1 |
|  | 40 | 100 | -14.9 | -4.5 | -19.4 |
|  | 155 | 81 | -0.2 | -1.2 | -1.4 |
|  | 156 | 94 | -16.6 | -5.0 | -21.6 |
|  | 341 | 100 | 0.1 | -5.3 | -5.2 |
|  | 343 | 100 | -48.9 | -6.1 | -55.0 |
|  | 370 | 100 | -17.7 | -4.9 | -22.6 |
|  | 371 | 100 | 5.3 | -8.6 | -3.3 |
|  | 372 | 100 | -16.7 | -14.4 | -31.2 |
|  | 373 | 100 | -17.5 | -1.8 | -19.3 |
|  | 394 | 100 | -44.4 | -1.3 | -45.6 |
| Pamidronate | 38 | 98 | -0.6 | -2.5 | -3.1 |
|  | 39 | 100 | -7.0 | -4.8 | -11.7 |
|  | 40 | 100 | -6.6 | -6.9 | -13.5 |
|  | 149 | 97 | -0.7 | -1.3 | -2.0 |
|  | 156 | 100 | -3.4 | -4.9 | -8.2 |
|  | 164 | 89 | -1.0 | -2.2 | -3.2 |
|  | 208 | 100 | 0.1 | -3.4 | -3.3 |
|  | 341 | 100 | 0.0 | -9.6 | -9.6 |
|  | 343 | 82 | -0.5 | -3.5 | -4.0 |
|  | 370 | 100 | -4.0 | -6.9 | -10.9 |
|  | 371 | 100 | 3.8 | -18.8 | -15.0 |
|  | 372 | 100 | -31.7 | -8.6 | -40.3 |
|  | 373 | 100 | -3.4 | -4.2 | -7.7 |
|  | 394 | 100 | -17.7 | -5.7 | -23.5 |

#### 3.6.4. Free Energy Landscape

The projections of the first (PC1) and second (PC2) eigenvectors were used to calculate the Free Energy Landscape (FEL) (Fig.6), with a color code where deeper blue color indicate wells of low free energy, where the protein conformation is more stable. The complexes *Tc*ENO-PEP (Fig.6b) and *Tc*ENO-etidronate (Fig.6d) had the most metastable conformational states, with local free energy wells distributed in four to five regions during the 50 ns simulation, while apo *Tc*ENO (Fig.6a) and *Tc*ENO-pamidronate (Fig.6c) had just two to three metastable conformations. In all the simulations the low energy configurations were explored several times by the protein. This results further confirm that binding to etidronate and PEP stabilizes the protein, while the complex with pamidronate has a stability comparable with the apo conformation.

**Fig. 6.**
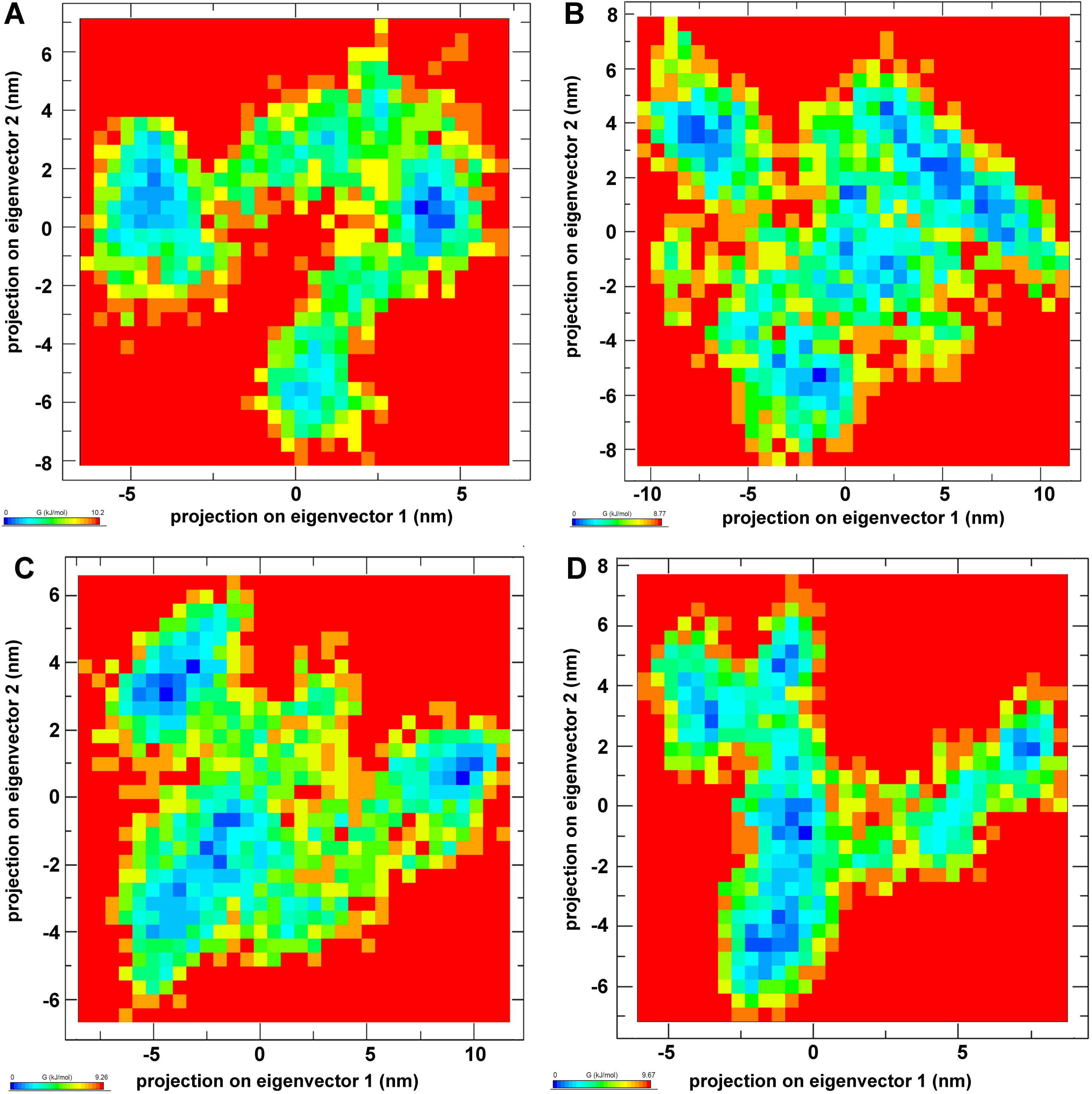
Free energy landscape (FEL) obtained from the first two eigenvectors of the MD simulated *Tc*ENO unbound (a), and in complex with PEP (b), pamidronate (c), and etidronate (d)

## 4. Conclusions

Enolase has been identified as a promising target in the search of drugs to treat Chagas disease, as well as other trypanosomatid caused diseases, for the essential role it plays in the energetic metabolism of the parasite and processes of transition between forms in its life cycle. In this work, we use previously characterized ligands as templates in the search of new possible inhibitors by similarity screening in a database comprising many approved drugs for its use in humans. After discarding metabolites retrieved in the search, a total of 6 compounds were used for molecular docking assays in order to predict their ability of binding to the active site of *Tc*ENO, followed by rescoring of the obtained poses. Only etidronate and pamidronate were predicted to be good binders to the enzyme, and were aligned with the known inhibitors to produce a pharmacophore model. Their binding modes and affinities were further explored in MD calculations, where only etidronate maintained the interactions predicted in the docking simulations, besides providing more stability to the enzyme in comparison with the apo conformation or in complex with pamidronate. Altogether, the results of *in silico* studies described here bestow molecular information about the *Tc*ENO binding, exploitable in the search and design of new drugs capable of inhibit the enzyme.

## Author contributions

EVV and CAP wrote the main manuscript text. EVV and CAP performed the computational simulations. MS, EVV, CR, CAP and MRM performed the data analysis. MS, EVV, CR, CAP and MRM reviewed the manuscript.

## Acknowledgments

We would like to thank Centro de Cómputos de Alto Rendimiento (CeCAR) for granting use of computational resources which allowed us to perform most of the MD calculations included in this work. A special thanks to Thomas Lemker from “Inte:Ligand Software Development & Consulting” for his help with the license of the LigandScout, and to Dr. Fernan Aguero for his help with the TDR Targets. This work was supported by Consejo Nacional de Investigaciones Científicas y Técnicas, Agencia Nacional de Promoción Científica y Tecnológica (FONCyT PICT 2015-0539). The research leading to these results has, in part, received funding from UK Research and Innovation via the Global Challenges Research Fund under grant agreement ‘A Global Network for Neglected Tropical Diseases’ grant number MR/P027989/1. CAP, MRM are members of the career of scientific investigator; EVV, MS and CR are research fellows from CONICET.

---

**Supplementary Figure 1.**
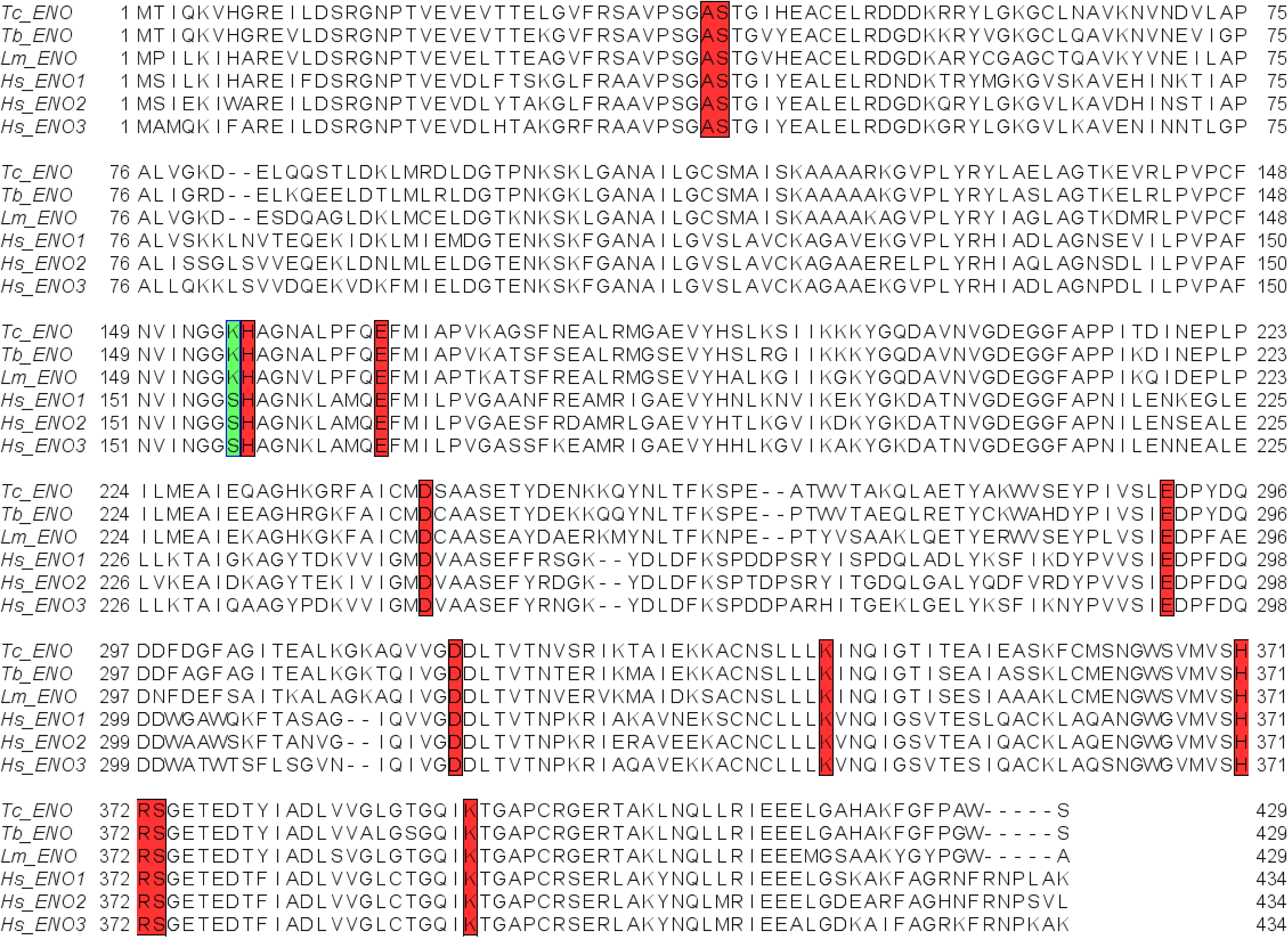
Amino acid multiple alignment of the enolases from *T. cruzi, T. brucei, L. mexicana*, and the three human enolases (ENO1, ENO2, and ENO3).

**Supplementary Figure 2.**
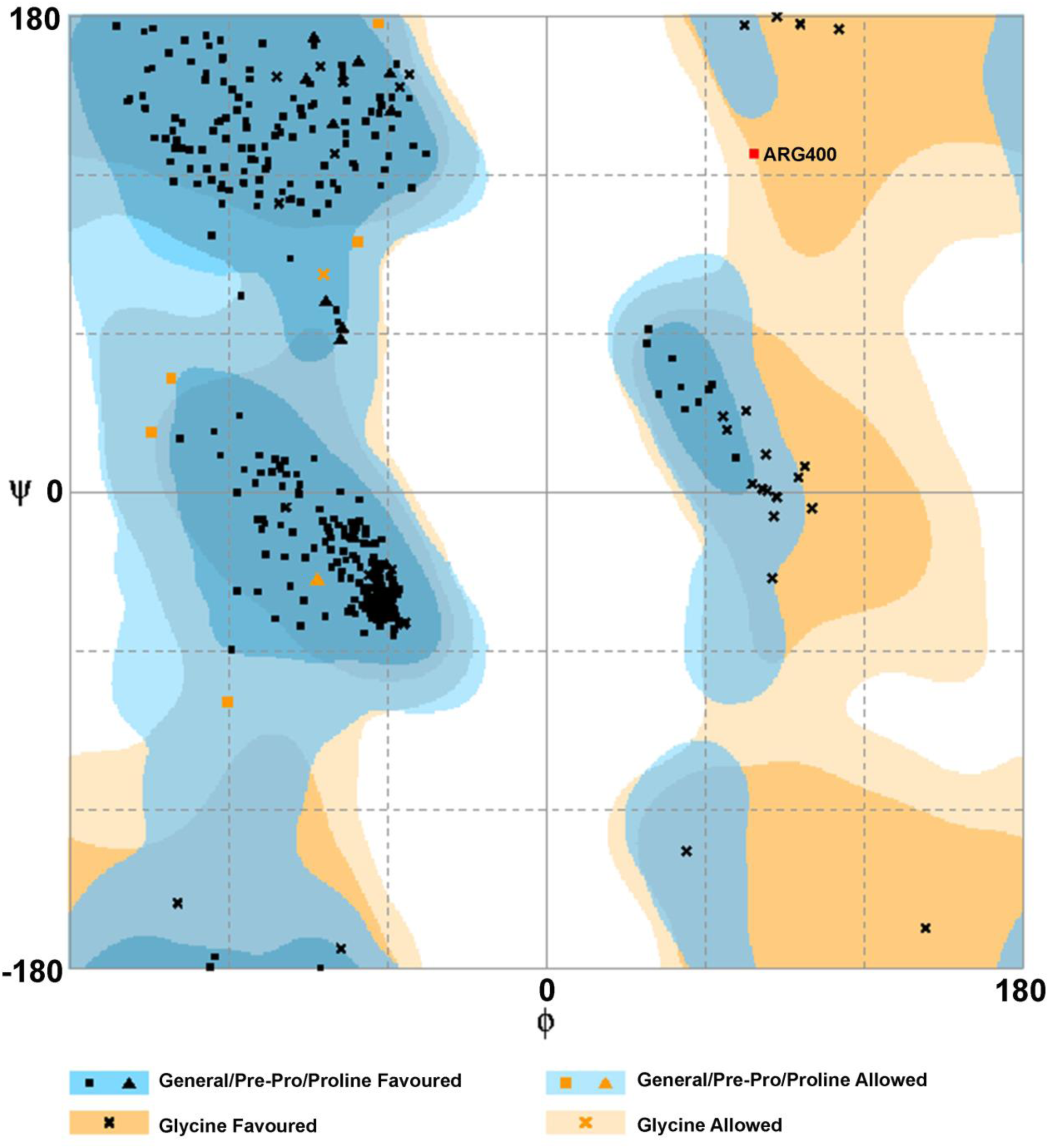
Ramachandran plot of the *Tc*ENO modelled with I-TASSER.

**Supplementary Figure 3.**
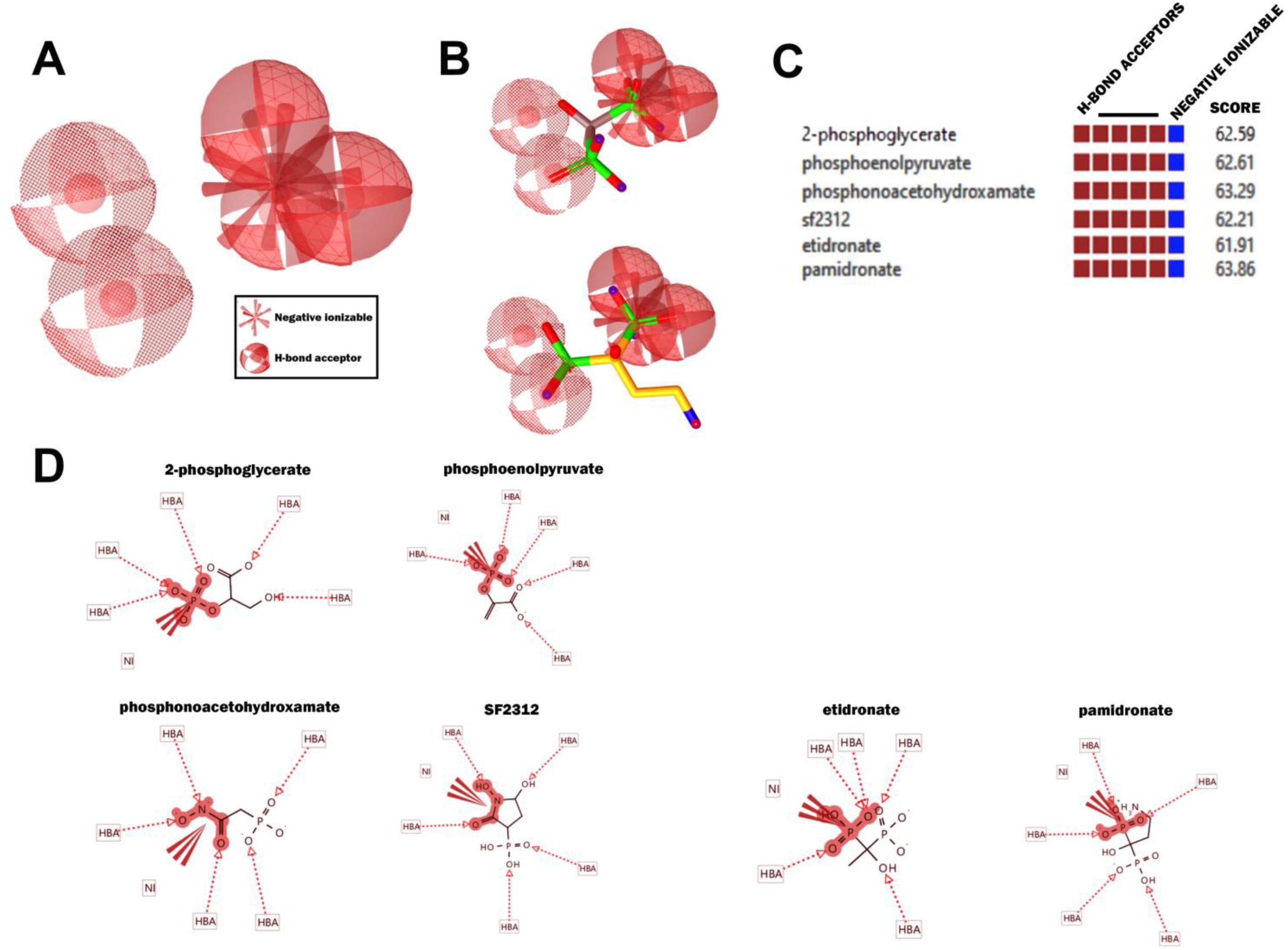
Pharmacophore mapping. A) Pharmacophore designed using the structures in common of the experimental binders of enolase and those predicted to bind in the molecular docking. Five H-bond acceptors are indicated as balls and a negative ionizable region as an asterisk (see reference). B) The structure of etidronate and pamidronate inside the pharmacophore representation. C) Table comparing the pharmacophore structures shared by the different compounds. D) Structures of the different compounds indicating the H-bond acceptors (HBA) and the negative ionizable groups (NI).

